# miR-2 contributes to WSSV infection by targeting Caspase 2 in mud crab (*Scylla paramamosain*)

**DOI:** 10.1101/2021.05.17.444593

**Authors:** Yi Gong, Jiao Chen, Yalei Cui, Shengkang Li

## Abstract

As we known, Caspase 2 is widely studied for its apoptosis regulatory function in mammals. However, despite the fundamental role of apoptosis during the anti-viral immune response, the relationship between Caspase 2 and virus infection has not been extensively explored in invertebrates, whether miRNAs are involved in this process also remains unclear. To address this issue, the miRNA-mediated regulation of Caspase 2 in mud crab *Scylla paramamosain* was characterized in this study. The results suggested that *Sp*-Caspase 2 could suppress white spot syndrome virus (WSSV) infection via apoptosis induction. The further data showed that Caspase 2 was directly targeted by miR-2 in mud crab. Silencing or overexpression of miR-2 could affect apoptosis and WSSV replication through regulating the expression level of Caspase 2. Taken together, all these results demonstrated the crucial role of miR-2-Caspase 2 pathway in the innate immunity of mud crab and revealed a novel mechanism during anti-viral immune response in marine invertebrates.

## Introduction

It is well known that crustaceans lack adaptive immunity and mainly rely on innate immune system (including humoral and cellular immunity) to recognize and protect the host against harmful microbes [1, 2]. In the recent years, growing evidence have demonstrated that the alterations associated with cell survival contribute to the pathogenesis of many diseases, including viral infections and autoimmune diseases [3]. Apoptosis, which is also called cell programmed death with specific features, including cytoplasmic narrow, membrane blebbing and chromatin aggregation [4], can be activated by various physiological and pathological stimulation [5]. It has been reported that virus infection would cause apoptosis activation in host [6], which could also serve as an effective defensive approach of the host cells to suppress virus invasion [7]. However, at present, the involvement of apoptosis during anti-viral immune response have not been extensively explored in marine invertebrates.

As we know, the caspase members are required for apoptosis induction and signal amplification [8]. Among them, as the second found caspase member, Caspase 2 is the most evolutionarily conserved in animals [9]. The gene for Caspase 2 was initially termed Nedd2 (neuronally expressed developmentally downregulated gene 2) during hybridization screening for mouse genes in neural precursor cells [10]. After that, Nedd2 was shown to be homologous to the *C. elegans* death gene-3 (CED-3) [11]. It has been reported that Caspase 2 is located in Golgi, mitochondria, nuclei and cytoplasm, and could serve as the upstream of DNA damage-induced apoptosis pathway by promoting cytochrome c release from mitochondria [12]. Moreover, cytotoxic stress could cause activation of Caspase 2, which is required for the permeabilization of mitochondria [13]. So far, Caspase 2-associated researches are mainly focused on higher animals, besides, the correlation with antiviral immune regulation also has not been addressed.

MicroRNAs (miRNAs), approximately 21∼27 nucleotide (nt) in length [14], are endogenous small non-coding RNA molecules that could bind to the 3’UTRs of target mRNAs by seed sequence complementary pairing, which further downregulate the expressions of target genes through translation repression or direct mRNA degradation [15]. It has been reported that miR-383 could promote human epithelial ovarian cancer development by inhibiting the expression of Caspase 2 [16]. Additionally, miR-708 could act as an oncogene and induce the carcinogenicity of bladder cancer by down-regulating Caspase 2 level [17]. Thus, it can be concluded that Caspase 2 was able to be regulated by miRNAs in various biological process. However, most of the researches associated with Caspase 2 targeted by miRNAs are mainly focused on cancer at present, whether Caspase 2 can be regulated by miRNAs and its roles has not been intensively investigated in invertebrates. To address this issue, the miRNA targeting Caspase 2 in mud crab was characterized in our study. White spot syndrome virus (WSSV), a large enveloped double-stranded DNA viral pathogen for many marine crustaceans [18], was used as an infection model. The results of this study revealed that miR-2 could affect apoptosis and WSSV replication through regulating the expression level of Caspase 2, indicating the crucial role of miR-2-Caspase2 pathway during anti-viral immune response of mud crab.

## Results

### Bioinformatics analysis of *Sp*-Caspase 2 cDNA sequence

The complete cDNA sequence of *Sp*-Caspase 2 contains an open reading frame (ORF) of 969 bp encoding 322 deduced amino acids, the sequence has been deposited in GenBank under the accession number MH558571.1. The putative protein sequence of *Sp*-Caspase 2 possesses a conserved CASc domain (amino acids 72-320) (Fig. 1A). Besides, the tertiary structure of *Sp*-Caspase 2 was predicted and the active sites were marked (Fig. 1B). Besides, the amino acid sequence of *Sp-*Caspase 2 was aligned with other crustaceans, the results showed that *Sp*-Caspase2 display the highest identity with *Pt*-Caspase 2 (92%) from *P. trituberculatus* (ARO92228.1) (Fig. 1C). Moreover, multiple sequence alignment indicated that the CASc domain of Caspase 2 was conserved across the species (Fig. 1C).

**Fig 1.**
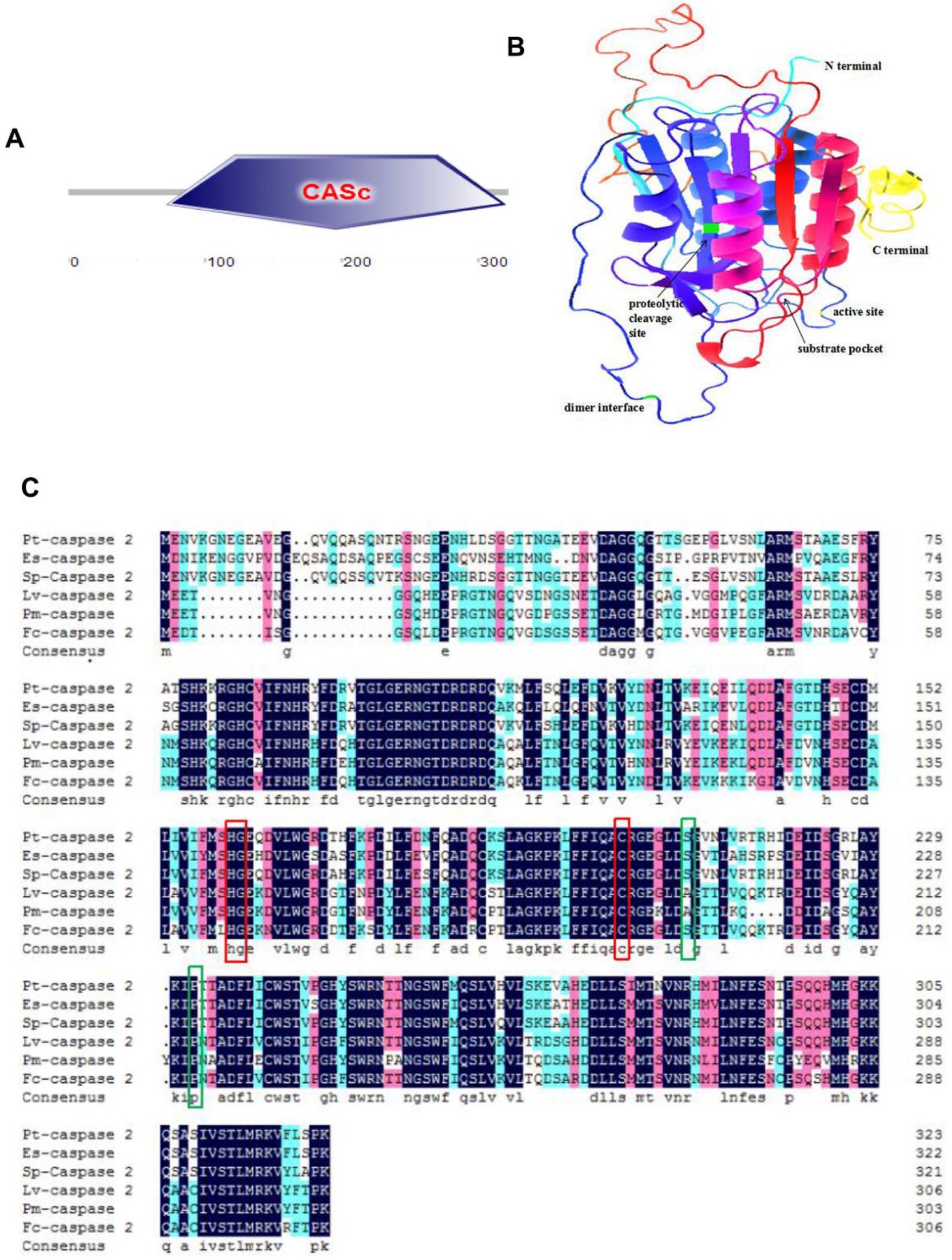
Bioinformatics analysis of *Sp-*Caspase 2. **(A)** Schematic view of the structure of Caspase 2 protein. **(B)** The three-dimensional model of Caspase 2 protein. **(C)** Multiple alignments of the deduced amino acid sequence of Caspase 2 protein in mud crab and other marine crustaceans. Conserved active sites (H158, C200) were marked with the red box, proteolytic cleavage sites (S207, P230) were marked with the green box. Proteins analyzed listed below: Pt-Caspase 2 (ARO92228.1) from *Portunus. trituberculatus*, Es-Caspase 2 (AGT29867.1) from *Eriocheir sinensis*, Lv-Caspase 2 (AGL61581.1) from *Litopenaeus vannamei*, Pm-Caspase 2 (ABO38430.1) from *Penaeus monodon* and Fc-Caspase 2 from *Fenneropenaeus chinensis* (ALL27850.1).

### Role of Caspase2 on virus infection in mud crab

To examine the effect of Caspase 2 on virus infection, we detected the expression level of Caspase 2 in mud crab challenged with WSSV. The results revealed that both mRNA and protein levels of Caspase 2 were upregulated during WSSV infection (Fig.2A and 2B), indicating that Caspase 2 may participate in the immune response to virus. To further ascertain whether Caspase 2 could affect WSSV proliferation in mud crab, Caspase 2 was silenced and then WSSV replication was evaluated. The results showed that the protein level of Caspase 2 was extremely decreased in mud crab after treated with Caspase 2-siRNA compared to control group (Fig. 2C). Besides, it was found that silence of Caspase 2-siRNA could suppress the expression of viral gene VP28 during WSSV infection (Fig. 2D). Taken together, these data strongly suggested that Caspase 2 is involved in immune response to virus infection, and could suppress virus proliferation in mud crab.

**Fig 2.**
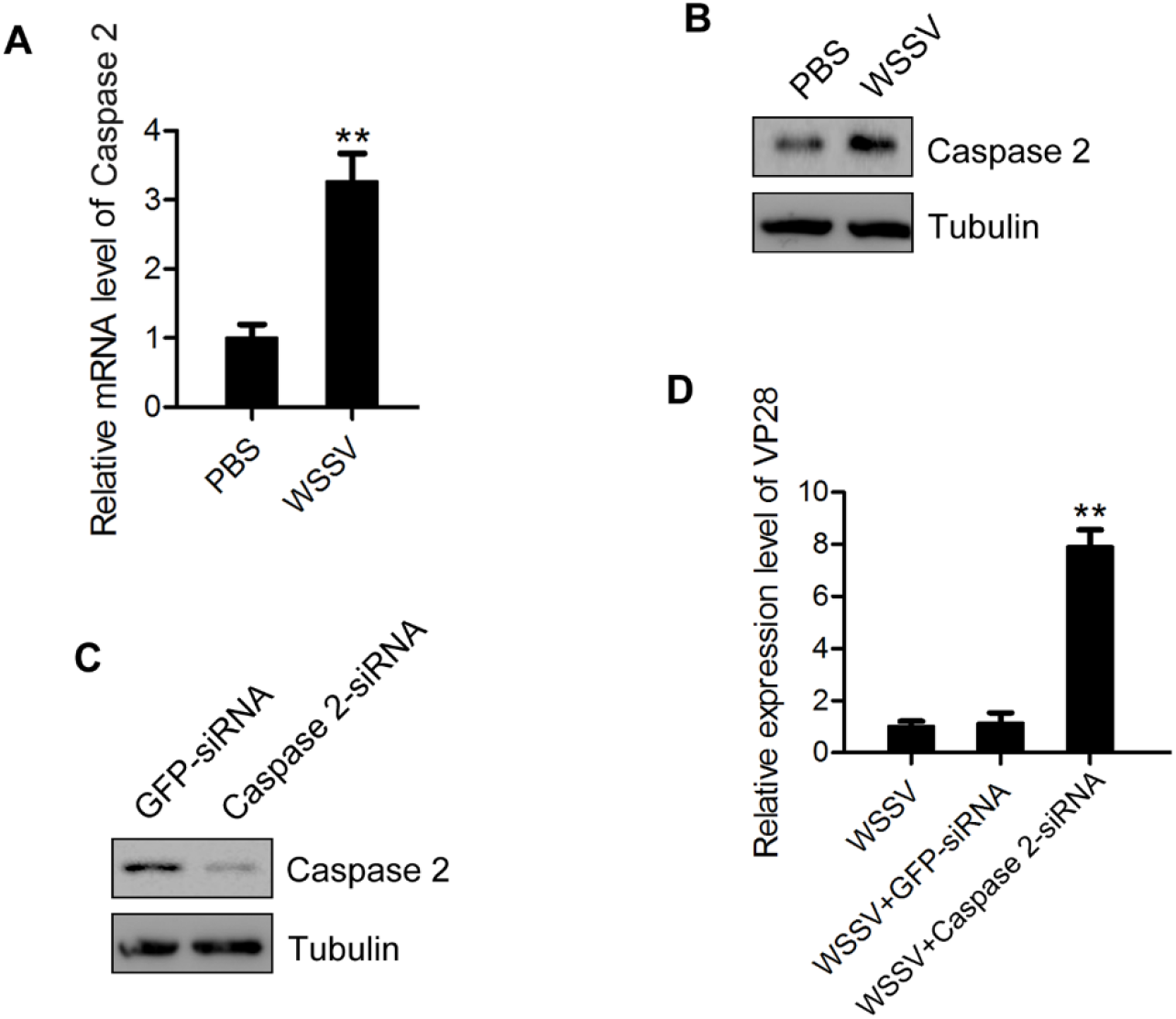
Role of Caspase 2 on virus infection in mud crab. **(A)** mRNA levels of Caspase 2 during virus infection in mud crabs. β-actin was used as an internal control. **(B)** Protein levels of Caspase 2 in the hemocytes of WSSV challenged crabs. Tubulin was used as an internal control. **(C)** The efficiency of Caspase 2 silence. Mud crabs were treated with Caspase 2-siRNA or Caspase 2-siRNA-scrambled, at 48 h post-injection, the Caspase 2 protein of hemocytes was detected by western blot. **(D)** The influence of Caspase 2 knockdown on WSSV infection in mud crab. Mud crabs were co-injected WSSV and Caspase 2-siRNA, followed by the expression detection of viral gene VP28 by qPCR. All data were the average from at least three independent experiments, mean ± s.d. (**, p<0.01).

### Effects of Caspase 2 in regulating apoptosis of mud crab

To further explore how Caspase 2 participate in the resistance to WSSV infection in mud crab, we attempt to detect Caspase 3/7 activity and apoptosis rate in the mud crabs treated with either WSSV or Caspase 2-siRNA. The results showed that both Caspase 3/7 activity and apoptotic rate of mud crab hemocytes were significantly increased following WSSV challenge compared with the control (Fig. 3A and 3B), indicating that virus infection could induce apoptosis in mud crab. Moreover, we found that the Caspase 3/7 activity and apoptotic rate of hemocytes were all decreased in mud crabs treated with WSSV and Caspase 2-siRNA compared to such in mud crabs treated with WSSV only (Fig. 3A and 3B). All these results demonstrated that Caspase 2 could protect mud crab from WSSV infection via promoting apoptosis.

**Fig 3.**
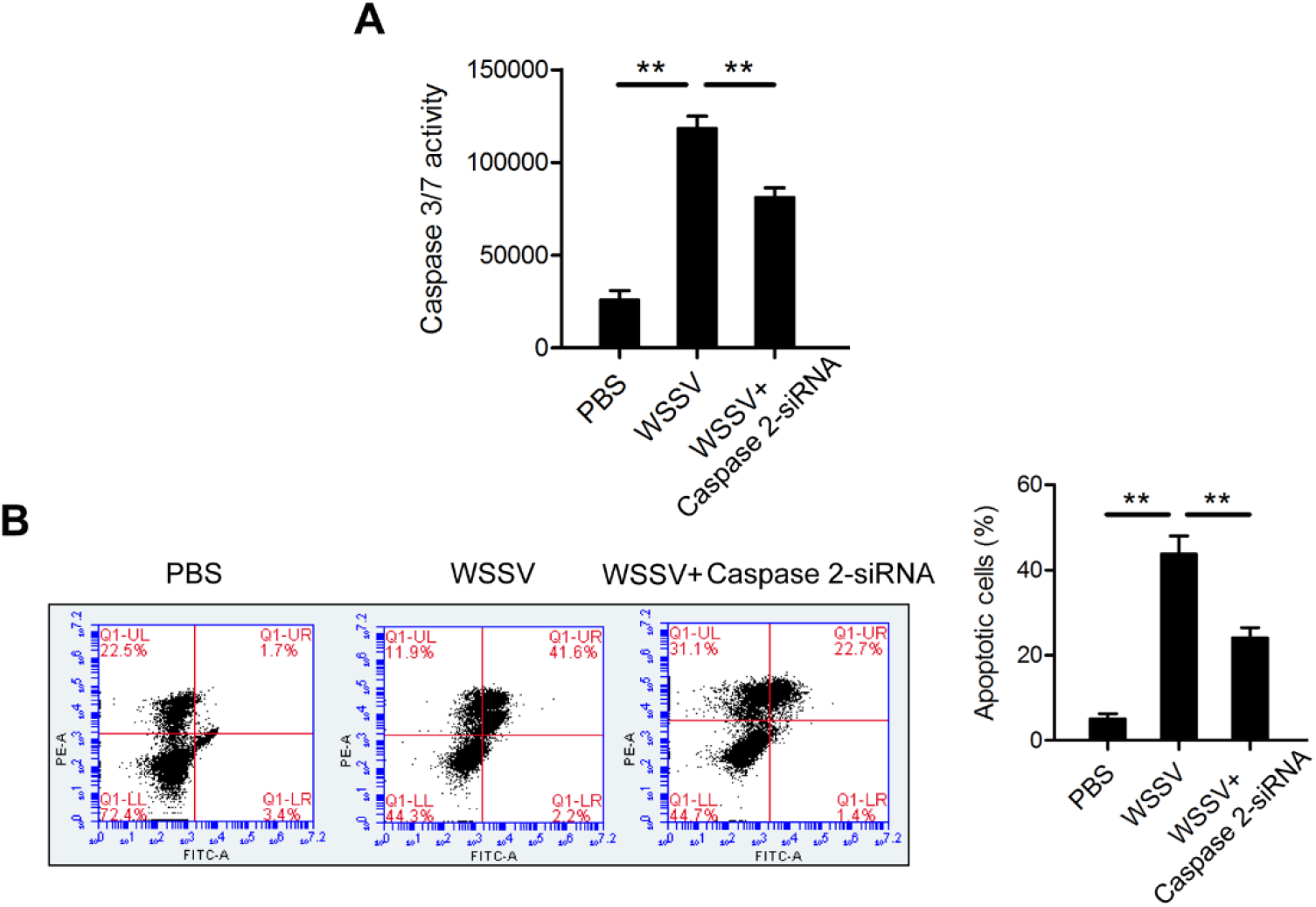
The function of Caspase 2 in regulating apoptosis. **(A-B)** The effect of Caspase 2 on apoptosis regulation of mud crab hemocytes. Mud crabs were injected with WSSV or co-injected with WSSV and Caspase 2-siRNA, then. the apoptotic levels of mud crab hemocytes were examined through the Caspase 3/7 activity detection **(A)** and annexin V analysis **(B)**. Data was shown as mean values ± standard deviations. Asterisks indicated significant differences (*P<0.05 and **P<0.01).

### The interactions between Caspase 2 and miR-2 in mud crab

To reveal the mechanism of Caspase 2 upregulation during virus infection in mud crab, miRNA targeting Caspase 2 were predicted using bioinformatics. The results showed that Caspase 2 could be potentially targeted by miR-2 (Fig. 4A). Then, to characterize the interaction between Caspase 2 and miR-2, the plasmid EGFP-Caspase consisting of EGFP and Caspase 2 3’UTR was constructed, and EGFP-Δ Caspase 2 3’UTR was served as control (Fig. 4B). After that, miR-2 and the constructed plasmids were co-transfected into S2 cells, the results showed that the fluorescence intensity of S2 cells treated with the EGFP-Caspase 2 3’UTR and miR-2 was significantly decreased compared with the controls, suggesting that miR-2 could bind with Caspase 2 3’UTR and inhibit its expression (Fig. 4C). Furthermore, we detected the subcellular location of Caspase 2 mRNA and miR-2 in the hemocytes of mud crabs by fluorescence in situ hybridization, the results indicated that miR-2 was co-localized with Caspase 2 mRNA (Fig. 4D).

**Fig 4.**
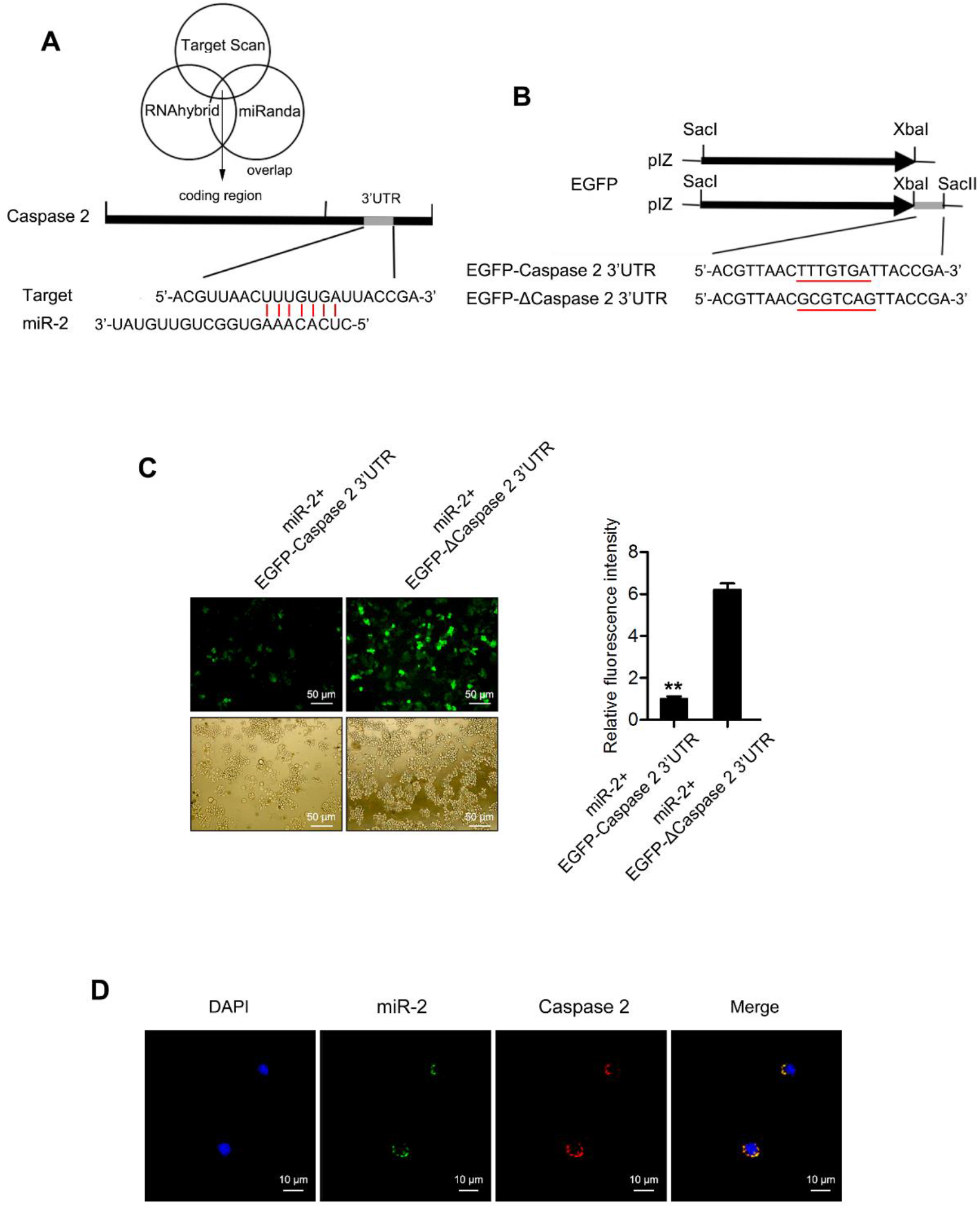
The interactions between miR-2 and Caspase 2 in mud crab. **(A)** The prediction of miRNA that targeting Caspase 2. Target Scan, RNAhybrid and miRanda algorithms were used for the prediction analysis, as predicted, miR-2 could target the 3’UTR of Caspase 2. **(B)** The construction of the plasmid EGFP-Caspase 2 3’UTR or EGFP-Δ Caspase 2 3’UTR. The seed sequence targeted by miR-2 was underlined. **(C)** The interaction between miR-2 and Caspase 2 3’UTR in S2 cells. S2 cells were co-transfected with miR-2 and the indicated constructed plasmids (EGFP-Caspase 2 3’UTR or EGFP-Δ Caspase 2 3’UTR), then the fluorescence intensity of S2 cells was detected and analyzed. **(D)** The co-localization of Caspase 2 mRNA and miR-2 in mud crab hemocytes. Caspase 2 mRNA probe was labeled with Cy3 (red), miR-2 probe was labeled with FAM (green), Scale bar, 10 μm. Significant statistical differences between treatment were indicated with asterisks (**, *p*<0.01).

To verify the effect of miR-2 on Caspase 2 expression, miR-2 was silenced or overexpressed in mud crabs, followed by detection of Caspase 2 levels. The results showed that both mRNA and protein levels of Caspase 2 were significantly increased when miR-2 was silenced in mud crab (Fig. 5A and 5B), while decreased in the miR-2 overexpressed mud crabs compared with the control group (Fig. 5A and 5B). Taken together, the above data suggested that miR-2 could directly interacted with Caspase 2 mRNA and suppress its expression.

**Fig 5.**
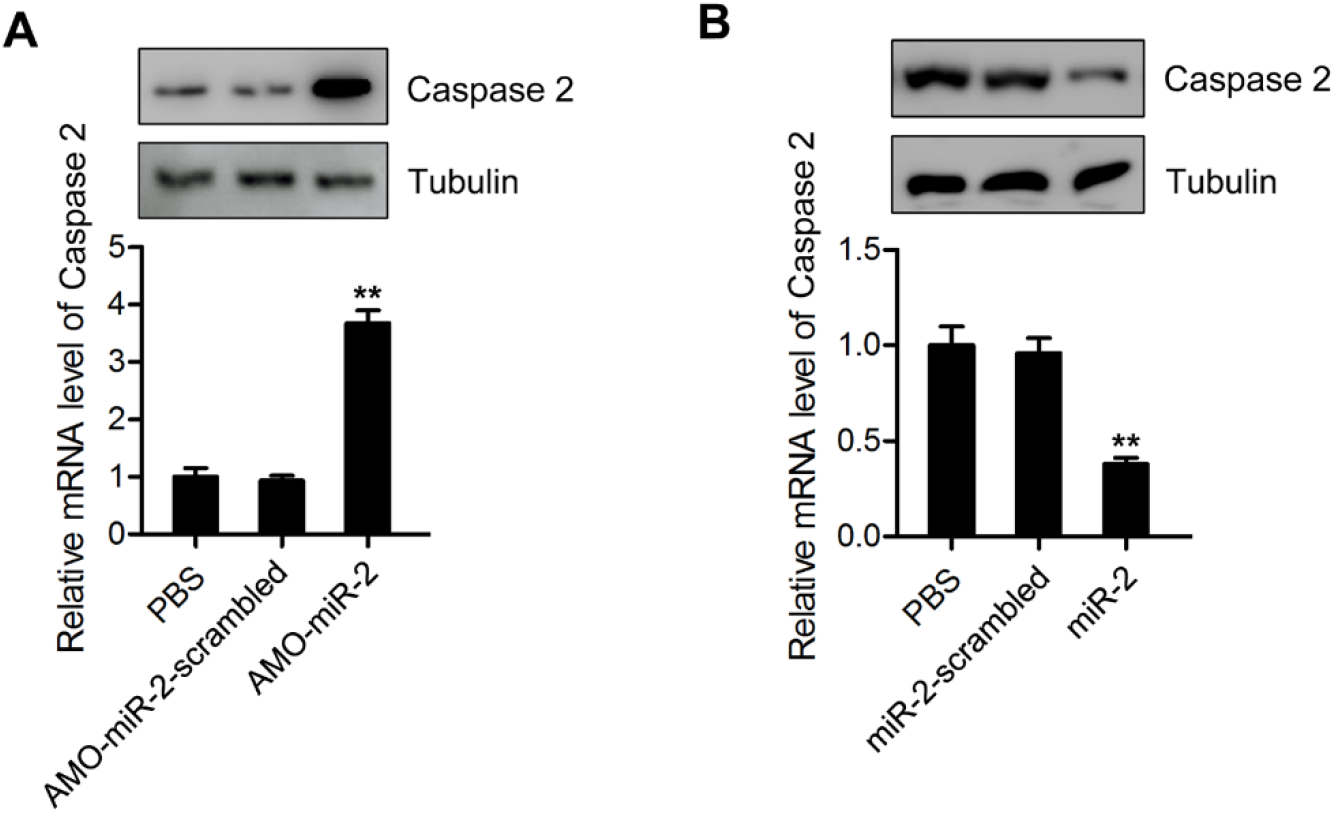
The effects of miR-2 on Caspase 2 expression in mud crab. **(A)** The influence of miR-2 silencing on the expression level of Caspase 2 in mud crabs injected with either AMO-miR-2 or AMO-miR-2-scrambled, and the mRNA and protein levels were detected at 48 h post-injection. **(B)** The influence of miR-2 overexpression on the expression level of Caspase 2 in mud crabs injected with either miR-2 or miR-2-scrambled, and the mRNA and protein levels were detected at 48 h post-injection. Each experiment was performed in triplicate and data are presented as mean ± s.d. (**, p<0.01).

### The involvement of miR-2-Caspase 2 pathway in response to WSSV infection

In an attempt to explore the role of miR-2 during virus infection, we detected the expression level of miR-2 during WSSV infection. The results of qPCR revealed that WSSV infection resulted in a significant decrease of miR-2 expression in mud crab (Fig. 6A), suggesting that miR-2 might participate in immune regulation of mud crab. To further evaluate the effect of miR-2 on virus infection, miR-2 was overexpressed or silenced during WSSV infection, the results showed that the expression of viral gene VP28 was significantly decreased in miR-2 silenced mud crab (Fig. 6B), while upregulated when miR-2 was overexpressed compared with the controls (Fig. 6C).

**Fig 6.**
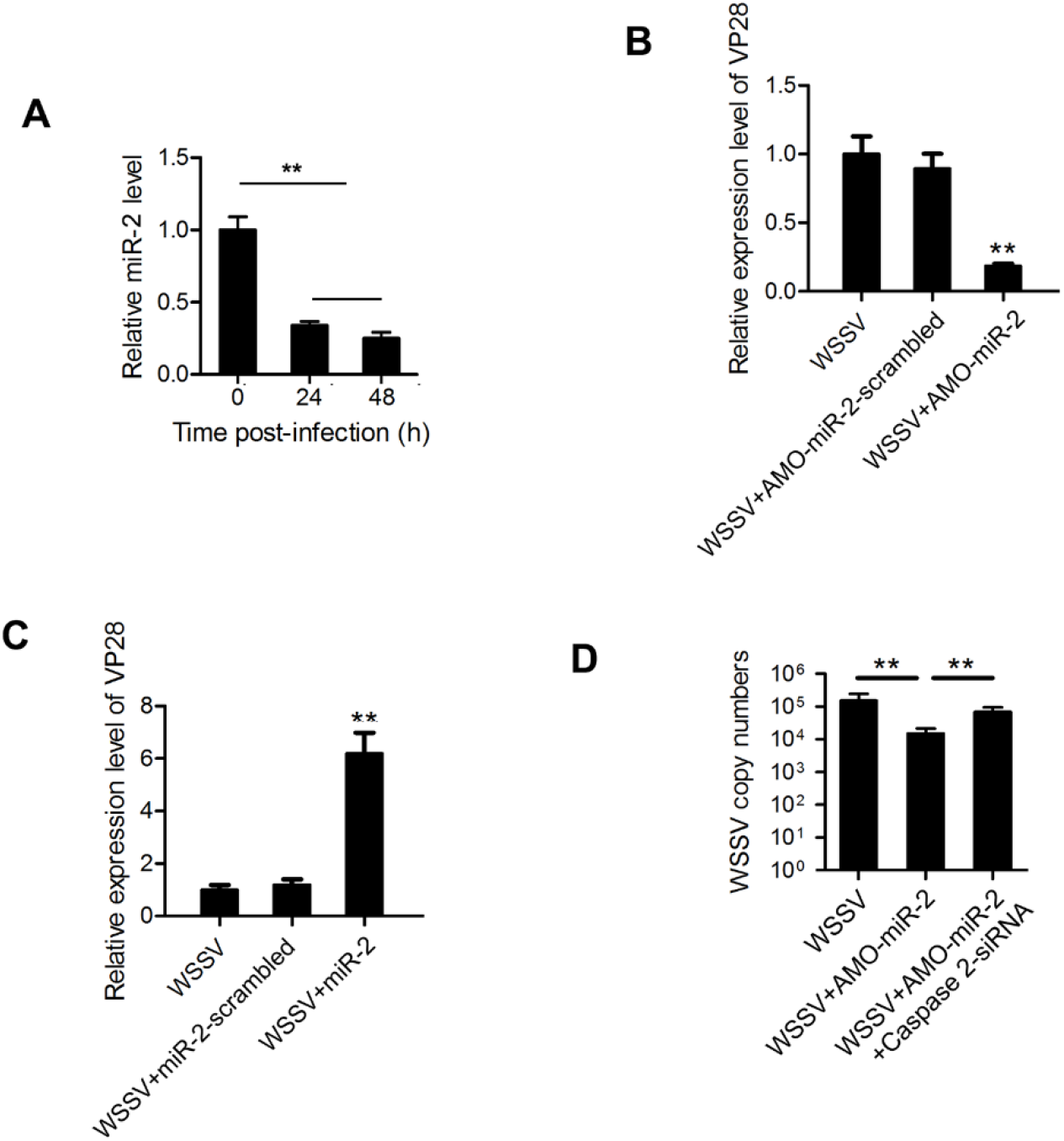
miR-2 promotes WSSV proliferation by targeting Caspase 2 in mud crab. **(A)** The detection of miR-2 expression during WSSV infection in mud crab using qPCR analysis. **(B)** The expression of virus gene VP28 in mud crabs co-treated with WSSV and AMO-miR-2, AMO-miR-2-scrambled was used as control. **(C)** The effects of miR-2 overexpression on the expression of virus gene VP28 in mud crab during WSSV infection. **(D)** The involvement of Caspase 2 during miR-2 -mediated virus promotion, AMO-miR-2, WSSV and Caspase 2-siRNA were co-injected into mud crabs, followed by the detection of WSSV copy numbers. All the numeral data represented the mean ± s.d. of triplicate assays (**, p<0.01).

Furthermore, when mud crab was co-treated with AMO-miR-2 and Caspase 2-siRNA, the AMO-miR-2-mediated virus suppression was remarkably relieved. These results revealed that miR-2 could promote WSSV infection by targeting Caspase 2 in mud crabs.

### The effects of miR-2-Caspase 2 pathway on apoptosis regulation

To explore the role of miR-2 during Caspase 2-mediated apoptosis regulation, the apoptosis activity was evaluated in mud crabs treated with AMO-miR-2 and Caspase 2-siRNA. The results showed that both Caspase 3/7 activity and apoptosis rate of mud crab hemocytes were significantly increased when miR-2 was silenced (Fig. 7A and 7B), suggesting that miR-2 was an anti-apoptotic miRNA in mud crab. Moreover, in the mud crab co-treated with AMO-miR-2 and Caspase 2-siRNA, the upregulated apoptosis activity in mud crab caused by miR-2 interference was significantly reduced, indicating that miR-2 could suppress apoptosis by targeting Caspase 2 in mud crab.

**Fig 7.**
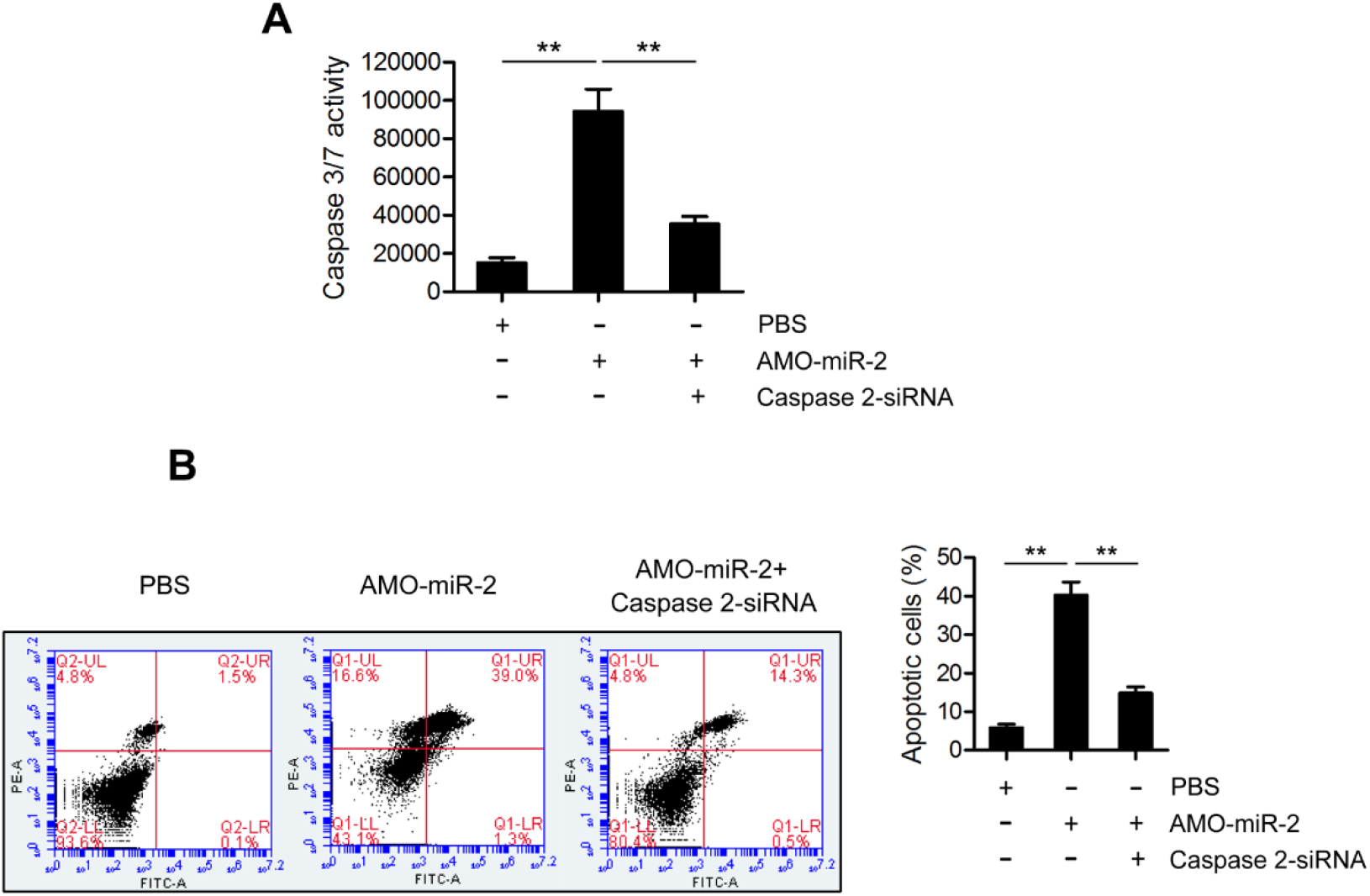
miR-2 suppresses apoptosis by targeting Caspase 2 in mud crab. **(A-B)** The involvement of Caspase 2 during the miR-2-mediated apoptosis regulation. Mud crabs were treated with either AMO-miR-2 or co-treated with AMO-miR-2 and Caspase 2-siRNA, followed by apoptosis evaluation in mud crab hemocytes through the Caspase 3/7 activity analysis **(A)** and annexin V assay **(B)**. Data presented were representatives of three independent experiments (**, *p*<0.01).

In summary, the above data showed that the expression of miR-2 was significantly decreased during WSSV infection in mud crab, resulting in the upregulation of Caspase 2 and the activation of cell apoptosis, which eventually suppressed virus replication in mud crab (Figure 8).

**Fig 8.**
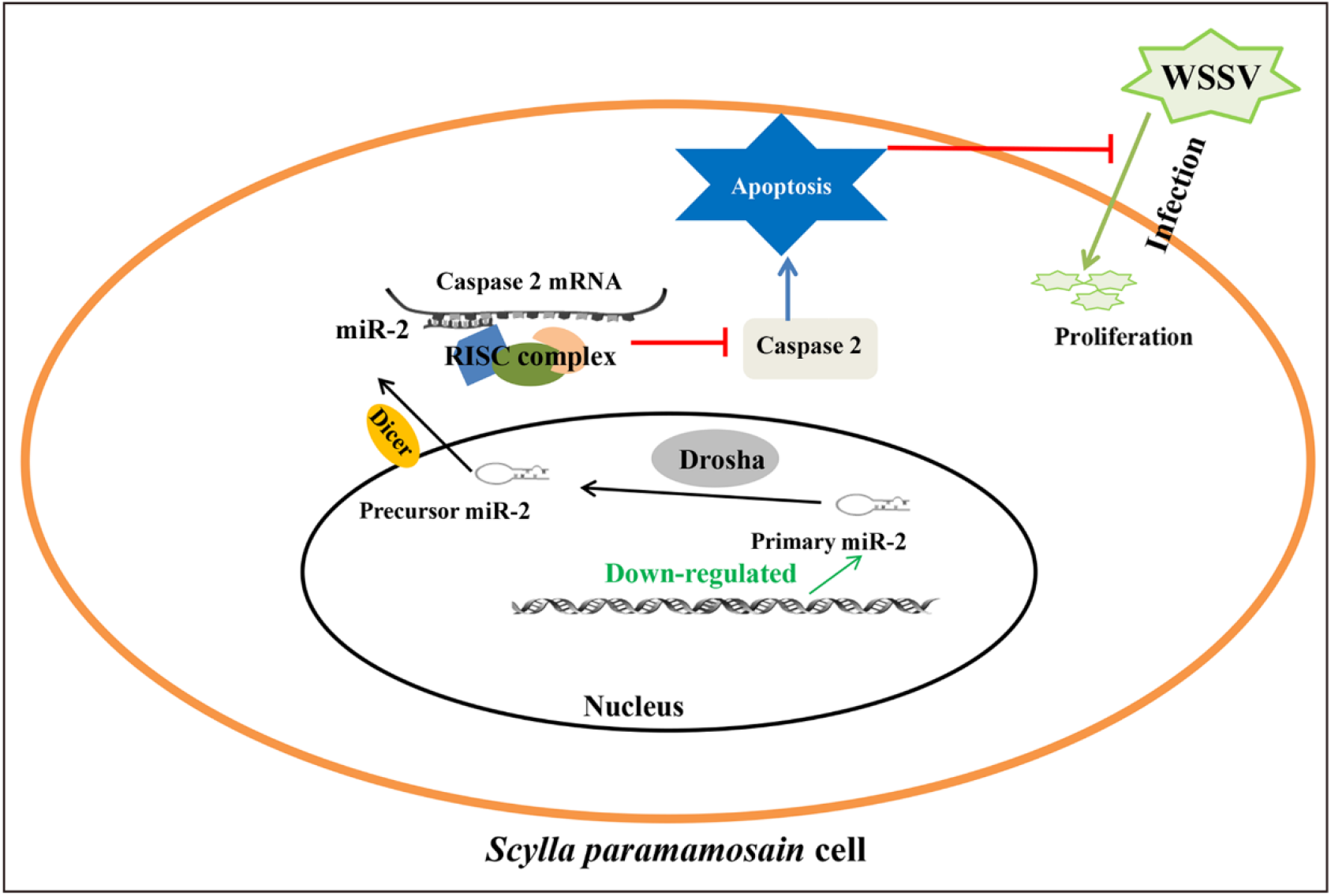
Proposed schematic diagram for miR-2-Caspase 2 pathway in regulating apoptosis and virus infection in mud crab.

## Discussion

In the past few years, it has become clear that the host could prevent virus replication and dissemination by apoptosis induction during infection process [19]. Shrimp miR-12 could simultaneously trigger phagocytosis and apoptosis of hemocytes through the synchronous downregulation of PTEN (phosphatase and tensin homolog) and TMBIM6 (transmembrane BAX inhibitor motif containing 6) in response to WSSV infection [20]. Similarly, through virus-host co-evolution, viruses also developed distinct strategies to overcome immunological defenses of the host by subverting host cell apoptosis [21]. It was found that the early-expressed nonstructural proteins (NS1 and NS2) of respiratory syncytial virus (RSV) could promote virus replication by delaying host cell apoptosis via IFN- and EGFR-independent pathway [22]. During WSSV infection, the virus-encoded miRNA WSSV-miR-N24 could target host caspase 8 gene and further repress apoptosis of shrimp hemocytes, which enhanced WSSV copies in shrimp [23]. At present, the relevant investigations performed in marine invertebrates are quite limited. In our study, we found that the apoptosis level in mud crabs was remarkably upregulated during WSSV infection, and demonstrated the essential role of miR-2-Caspase 2 pathway in the regulation of apoptosis and virus infection. Therefore, our findings revealed a novel miRNA-mediated regulatory mechanism during antiviral immune response in mud crab.

The first described feature of Caspase 2 is the activator of extrinsic apoptosis pathway [24]. The mitosis-promoting kinase, cdk1-cyclin B1 can phosphorylate Caspase 2 at Ser 340 to prevent its activation, which further suppresses apoptosis upstream of mitochondrial cytochrome c release [25]. In the recent years, Caspase-2 has also been proved to mediate non-apoptotic signaling pathways. It has been reported that Caspase 2 could serve as a tumor suppressor in Kras (kirsten rat sarcoma viral oncogene)-driven lung cancer, silencing of Caspase 2 would lead to enhanced tumor proliferation and progression [26]. Also, Caspase 2 was found to form complexes with the cell cycle regulatory proteins cyclin D3, CDK4, and p21/Cip1 to promote AR (androgen receptor) transactivation by inhibiting the repressive function of cyclin D3 [27]. So far, Caspase 2 has been widely studied for its apoptotic and non-apoptotic functions, however, the relationship between Caspase 2 and virus infection has not been previously addressed. In this study, the involvement of Caspase 2 during WSSV infection was determined, the results showed that both mRNA and protein levels of Caspase 2 in mud crab was significantly upregulated after WSSV challenge, and the copy numbers of WSSV was increased when Caspase 2 was silenced. For the first time, the present study revealed the crucial role of Caspase 2 in the immune response to virus infection in mud crab.

RNAi, mainly mediated by siRNAs or miRNAs, was a natural defensive approach to virus infection through post-transcriptional gene regulation [28]. It has been proved that the miRNA-mediated RNAi pathways were important regulators in many biological processes [29]. Silencing of miR-100 would result in the increase of apoptotic activity of shrimp hemocytes by regulating the expression of trypsin, which further led to the decreases of virus genome copies during WSSV infection [30]. Besides, it was found that miR-200c could be induced by oxidative stress and caused endothelial cell apoptosis and senescence by targeting ZEB1 (Zinc finger E-Box binding homeobox 1) [31]. Previous researches have demonstrated that miRNAs were tightly relevant to the regulation of apoptosis and immune regulation in both vertebrates and invertebrates. However, whether Caspase 2 could be regulated by miRNAs and their biological significances remains unaddressed in invertebrates. Here, we revealed the transcriptional crosstalk between miR-2 and Caspase 2 during WSSV infection in mud crab, and found that the miR-2 was downregulated after WSSV challenge, leading to the accumulation of Caspase 2, which eventually triggered apoptosis and attenuated WSSV replication in mud crab. In this context, our findings provided a clue to clarify the role of miRNAs during Caspase 2-mediated apoptosis regulation and virus suppression in marine invertebrates.

## Materials and methods

### Mud crab culture and WSSV challenge

Healthy mud crabs were purchased from Niutianyang farm (Shantou, Guangdong, China) and acclimated in tanks under laboratory conditions (10 ‰ salinity, 25 °C) for a week before experiments. Then, each crab was injected with 200 μL of WSSV solution (1×10^6^ copies/mL) according to our previous study [32], PBS-treated mud crabs was served as control group. At different times post-infection, hemocytes and muscles were collected from three randomly chosen crabs per group and stored at -80 °C for later use.

### RNA interference of Caspase 2

The siRNA targeting mud crab Caspase 2 was designed with BLOCK-iT RNAi Designer (https://rnaidesigner.lifetechnologies.com/rnaiexpress/sort.do) based on the sequence of Caspase 2 (GenBank accession number MH558571.1), generating Caspase2-siRNA (5’-UGUUACACGGUCAAAGUAGCGUU-3’) and its control Caspase2-siRNA-scrambled (5’-AAGAGCGAUGGCGUAUACUUCUU-3’). The siRNAs were synthesized using the *in vitro* transcription T7 kit (TaKaRa, Japan). Then, 50 μg of Caspase2-siRNA or Caspase2-siRNA-scrambled was injected into each mud crab in two doses, at intervals of 12 h. At different time post the last injection, three mud crab were randomly selected for each treatment at stored at -80 °C for later use.

### Analysis of WSSV copies with qPCR

The genomic DNA of WSSV was extracted from crab muscle with a SQ tissue DNA kit (Omega Bio-tek, USA) according to manufacturer’s instruction, the copy number of WSSV was detected by qPCR analysis using Premix Ex Taq (Probe qPCR) (TaKaRa, Japan). The qPCR was performed with WSSV-specific-primers (5’-TTGGTTTCATGCCCGAGATT-3’ and 5’-CCTTGGTCAGCCCCTTGA-3’) and a TaqMan probe (5’-FAM-TGCTGCCGTCTCCAA-TAMRA-3’) as described in our previous study [33].

### Quantification of mRNA with qPCR

Total RNAs were extracted from crab hemocytes by RNA isolation kit (Ambion, USA). Reverse transcription reaction was conducted with PrimeScript™ RT Reagent Kit (Takara, Japan). The primers (5’-GGGACAAGGAACAACAGAAT-3’ and 5’-ACACGGTCAAAGTAGCGATG-3’) were used to quantify the Caspase 2 mRNA, β-actin was served as the internal control, which quantified with primers (5’-GCGGCAGTGGTCATCTCCT-3’ and 5’-GCCCTTCCTCACGCTATCCT-3’). Then, the relative fold change of Caspase 2 expression levels was determined using the 2^−ΔΔCt^ algorithm.

### Bioinformatics analysis of Caspase2

The comparison of Caspase2 full-length cDNA and deduced amino acid sequences between different species was conducted by BLAST software at the National Center for Biotechnology Information (http://www.ncbi.nlm.nih.gov/blast/). The deduced amino acid was obtained by ORF Finder (http://www.ncbi.nlm.nih.gov/gorf/orfig.cgi), then Signal P 3.0 program was used to predict the presence and location of signal peptide (http://www.cbs.dtu.dk/services/SignalP). The transmembrane domain was predicted by the TMHMM (http://www.cbs.dtu.dk/services/TMHMM). Besides, the tertiary structure of Caspase 2 protein was designed online and edited by PyMol Viewer (https://zhanglab.ccmb.med.umich.edu/I-TASSER).

### Detection of apoptotic activity

To detect apoptosis rate, the hemocytes of crabs were collected by centrifugation at 600×g for 10 mins at 4 °C, then, the hemocytes were washed twice and treated according to protocol of BD Pharmingen FITC Annexin V Apoptosis Detection Kit (BD Biosciences, US). In addition, the hemocytes were also placed onto a 96-well plate at a density of 10^5^ hemocytes per well, after added 100 mL of Caspase-Glo 3/7 reagent (Promega, USA), the plate was gently shaken at room temperature for 2 h and the luminescence was measured by a plate-reading luminometer.

### Cell culture, transfection, and fluorescence assays

The Drosophila Schneider 2 (S2) cells (Invitrogen) were co-transfected with the EGFP-Caspase 2 or EGFP-Δ Caspase 2 plasmid and the synthesized miR-2 (miR-2-mimic) using Cellfectin II transfection reagent (Invitrogen) according to the manufacturer’s protocol. The concentrations of miR-2-mimic and the plasmid were 50 nM/well and 100 ng/well, respectively. At 48 h after co-transfection, the EGFP fluorescence intensity was measured by microplate reader at 490/ 510 nm of excitation/emission (Ex/Em).

### Fluorescence in situ hybridization

The hemocytes were seeded onto polysine-coated coverslips and then fixed with 4% polyformaldehyde for 15 min at room temperature. Followed by dehydrated in 70% ethanol at 4 °C overnight. After that, the coverslips were incubated with hybridization buffer [1× SSC (15 mM sodium citrate, 150 mM sodium chloride, pH 7.5), 10% (w/v) dextran sulfate, 25% (w/v) formamide, 1× Denhardt’s solution] containing 100 nM of probe at 37 °C for 5 h. The probe used is listed below, miR-2 probe (5’-FAM-ATACAACAGCCACTTTGTGAG-3’), Caspase 2 mRNA probe (5’-Cy3-TCCAGCA AGAGACTTGCACTGA-3’). Subsequently, the slips were washed with PBS three times and stained with DAPI (4’, 6-diamidino-2-phenylindole) (50 ng/mL) (Sigma, USA) for 5 min, and observed by CarlZeiss LSM710 system (Carl Zeiss, Germany).

### The silencing or overexpression of miR-2 in mud crab

The mimic of miR-2 (miR-2-mimic) or anti-miR-2 oligonucleotide (AMO-miR-2) was injected into crabs at 30 μg/crab to overexpress or knockdown miR-2 in mud crab. miR-2-mimic (5’-CUCACAAAGUGGCUGU**U**GU**A**U-3’) and AMO-miR-2 (5’-AUACAACAGCCACUUU**G**UG**A**G-3’) were all synthesized by Sangon Biotech (Shanghai, China) and modified with 2’-O-methyl (OME) (bold letters) and phosphorothioate (the remaining nucleotides). At different time post injection, three crabs from each treatment were randomly collected for later use.

### Quantification of miR-2 with qPCR

Total RNA of mud crab was extracted via MagMAX mirVana Total RNA Isolation Kit (Thermo Fisher Scientific, USA), then, the isolated RNAs were subjected to first-strand cDNA synthesis using (5’-GTCGTATCCAGTGCAGGGTCCGAGGTCACTGGATACGACATACAACA-3’) by PrimeScriptTM II 1st Strand cDNA Synthesis Kit (Takara, Japan). After that, Premix Ex Taq (TaKaRa, Japan) was used to quantify the expression level of miR-2, U6 was used as the internal control. The relevant primers used were listed below, miR-2 (5’-CGCCGCTCACAAAGTGGC-3’ and 5’-TGCAGGGTCCGAGGTCACTG-3’), U6 (5’-CTCGCTTCGGCAGCACA-3’ and 5’-AACGCTTCACGAATTTGCGT-3’).

### Statistical analysis

All the numerical data presented were analyzed by one-way analysis of variance (ANOVA) to calculate the means and standard deviations of triplicate assays. The differences were considered statistically significant at P<0.05 and P<0.01.

## Acknowledgments

This study was financially supported by the National Natural Science Foundation of China (31802341, 31902406), STU Scientific Research Foundation for Talents (NTF18001).

## Author contributions

YG and JC performed the experiments and analysed the data, YLC provided technical supports, YG and SKL wrote the manuscript. All authors read and approved the contents of the manuscript and its publication.

## Disclosure statement

The authors declare no conflicts of interest.

